# An essential function for autocrine Hedgehog signaling in epithelial proliferation and differentiation in the trachea

**DOI:** 10.1101/2022.01.13.476169

**Authors:** Wenguang Yin, Andreas Liontos, Janine Koepke, Maroua Ghoul, Luciana Mazzocchi, Xinyuan Liu, Chunyan Lu, Haoyu Wu, Athanasios Fysikopoulos, Alexandros Sountoulidis, Werner Seeger, Clemens Ruppert, Andreas Günther, Didier Y.R. Stainier, Christos Samakovlis

## Abstract

The tracheal epithelium is a primary target for pulmonary diseases as it provides a conduit for air flow between the environment and the lung lobes. The cellular and molecular mechanisms underlying airway epithelial cell proliferation and differentiation remain poorly understood. Hedgehog (Hh) signaling orchestrates communication between epithelial and mesenchymal cells in the lung, where it modulates stromal cell proliferation, differentiation and signaling back to the epithelium. Here, we reveal a new, autocrine function of Hh signaling in airway epithelial cells. Epithelial cell depletion of the ligand Sonic hedgehog (SHH) or its effector Smoothened (SMO) causes defects in both epithelial cell proliferation and differentiation. In cultured primary human airway epithelial cells, Hh signaling inhibition also hampers cell proliferation and differentiation. Epithelial Hh function is mediated, at least in part, through transcriptional activation as Hh signaling inhibition leads to downregulation of cell-type specific transcription factor genes in both the mouse trachea and human airway epithelial cells. These results provide new insights into the role of Hh signaling in epithelial cell proliferation and differentiation during airway development.

## INTRODUCTION

Hedgehog (Hh) signaling regulates tissue patterning and cell differentiation in most animals. In mammals, there are three Hh ligands, desert hedgehog (DHH), indian hedgehog (IHH) and sonic hedgehog (SHH). Hh signaling has been investigated in depth during limb formation (Riddle et al., 1993), endodermal organ development (Sala et al., 2011; Bellusci et al., 1997), and cell differentiation in the neural tube (Ericson et al., 1995).

In the lung epithelium, SHH is a highly expressed ligand coordinating both organ formation and homeostasis (Sala et al., 2011; Bellusci et al., 1997; Peng et al., 2015). Epithelial SHH activates the transmembrane protein Smoothened (SMO) in mesenchymal cells leading to nuclear translocation and activation of GLI transcription factors. The activation of Hh signaling in the signal-receiving cells depends on the negative regulators PTCH1 and HHIP, whose transcription is in turn activated by GLI proteins, creating a tightly regulated negative feedback loop (Wang et al., 2018; Kugler et al., 2015). During lung development, the activation balance of Gli1, Gli2 and Gli3 controls cell proliferation and differentiation of mesenchymal airway smooth muscle cells (Li et al., 2004) and endothelial cells (Li et al., 2004; Miller et al., 2004). In the mesenchyme, Hh signaling activation also controls the expression of genes encoding signals like FGF10 or BMP4 (Weaver et al., 2003), which become activated or restricted in highly localized domains to shape epithelial branching and morphogenesis (Pepicelli et al., 1998; Herriges et al., 2015). In adult lungs, Hh signaling is activated upon injury (Watkins et al., 2003) and recent studies have shown that activation of Hh signaling in Gli1^+^ mesenchymal cells maintains quiescence in club cells (Peng et al., 2015). In the alveolar compartment, asymmetric Hedgehog signaling activation in the distal mesenchymal cells marked by *Gli2* and *Pdgfrα* expression controls alveolar epithelial cell proliferation (Wang et al., 2018). Ectopic activation of Hh signaling in the distal mesenchyme leads to emphysema-like phenotypes (Wang et al., 2018). These results highlight a central function for Hh signaling in adult lung homeostasis. In parallel, several genome-wide association studies (GWAS) have identified polymorphisms in genes encoding Hh signaling components in cohorts of asthma and COPD patients implicating Hh signaling in chronic lung disease initiation or progression (Kugler et al., 2015; Wang et al., 2019). Overall, genetic studies of Hh signaling in the mouse embryonic and adult lungs have focused on the paracrine role of epithelial SHH in controlling the patterning of the mesenchyme. Mesenchymal cells respond to SHH by the restricted activation of distinct programs that in turn, guide epithelial morphogenesis and tissue repair responses in the adult. Although the paracrine functions of SHH are instrumental in the lung, its potential autocrine function in respiratory tract development has not been formally investigated. Recent pharmacological evidence has suggested a role of SHH in the proliferation and differentiation of cultured human primary nasal epithelial (HNE) cells (Belgacemi et al., 2020), but the role of epithelial Hh signaling in the lung remains elusive.

The tracheal tube consists of endoderm-derived epithelium surrounded by mesoderm-derived cartilage, connective tissue, and smooth muscle (Brand-Saberi et al., 2014). The tracheal epithelium is composed of several cell types including basal progenitor cells, ciliated cells, club cells, goblet cells, neuroendocrine cells, tuft cells and ionocytes (Montoro et al., 2019; Plasschaert et al., 2018). Like the lung airways, tracheal tube formation depends on complex epithelial*-*mesenchymal interactions (Sala et al., 2011; Hines et al., 2013; Snowball et al., 2015; Gerhardt et al., 2018) that orchestrate cellular proliferation (Snowball et al., 2015) and differentiation (Gerhardt et al., 2018). During embryonic development, inactivation of Hh signaling impairs tracheal cartilage formation by reducing SOX9^+^ chondrocyte proliferation (Park et al., 2010), differentiation (Park et al., 2010) and condensation (Yin et al., 2018). This process involves the interplay of SHH emanating from the tracheal epithelium and FGF10 from the mesenchyme (Sala et al., 2011). A recent genetic study in pediatric patients associated Hh signaling with tracheal ring deformity characterized by complete cartilaginous rings surrounding the trachea and tracheal stenosis (Sinner et al., 2019).

Here, we investigate the potential role of Hh signaling in tracheal epithelial cells. We first show that Hh signaling components are dynamically expressed in the tracheal epithelium and mesenchyme. Conditional *Shh* inactivation in *Nkx2*.*1*-expressing epithelial cells led to early postnatal lethality accompanied by respiratory defects. The tracheal cartilage rings were discontinuous and the tracheal tube collapsed. Epithelial cell proliferation and the number of differentiated secretory and multiciliated cells were reduced in the *Shh* mutant trachea. Inactivation of the transmembrane protein *Smo* or expression of a dominant active form of *Smo* in the lung endoderm caused opposite defects in tracheal epithelial cell proliferation and differentiation arguing for an autocrine role of Hh signaling in directly promoting proliferation and differentiation in the tracheal epithelium. We used an *in vitro* air-liquid interface (ALI) differentiation model of primary human bronchial cells to test whether the autocrine role of Hh signaling functions in isolated human epithelial cells. Chemical inhibition of SMO, or GLI, led to defects in epithelial cell proliferation and differentiation arguing for an epithelial cell-autonomous function of Hh signaling in human bronchial cells as well. Our results reveal a new autocrine function of Hh signaling in addition to its previously characterized paracrine roles in respiratory organ development and prompt further investigations of epithelial Smo-mediated Hh signaling in lung epithelial cells during embryonic development and lung disease progression.

## RESULTS

### Epithelial deletion of Shh results in tracheal tube formation defects

To examine the spatiotemporal mRNA expression of Hh signaling components in the developing trachea, we performed multiplex fluorescence *in situ* hybridizations using SCRINSHOT (Single Cell Resolution IN Situ Hybridization On Tissues) during embryonic stages (Sountoulidis et al., 2020). We examined the cellular colocalization of *Shh, Smo, Patched 1* (*Ptch1*), *Patched 2* (*Ptch2*), *Hedgehog interacting protein* (*Hhip*), *Gli1, Gli2* and *Gli3* mRNAs in epithelial and mesenchymal cells from E13.5 to E15.5 (Fig. 1a-b). To mark epithelial cells, we used probes against the *Cdh1* and *Trp63* mRNAs. *Shh* was expressed in the epithelium and *Hhip* in the mesenchyme and both displayed reduction in expression levels from E13.5 to E15.5 (Fig. 1a, b). The Hh signaling transducer gene *Smo* was expressed in both epithelial and mesenchymal cells including putative chondroblasts (Fig. 1a, b and Supplementary Fig. 1). Similarly, the Hh signaling transcription factor genes (*Gli1, Gli2 and Gli3*) and transmembrane receptor genes (*Ptch1* and *Ptch2*) were detected in both epithelial and mesenchymal cells and displayed higher expression levels at E13.5 compared to E15.5 (Fig. 1a, b). We also examined *Shh, Smo, Gli1* and *Gli2* mRNA levels in whole tracheal extracts by quantitative reverse transcription PCR (RT-qPCR). *Shh, Smo, Gli1* and *Gli2* mRNAs were detectable as early as E12.5 and became gradually reduced until E18.5 (Supplementary Fig. 2a-d). *Smo* mRNA levels did not dramatically change possibly due to mesenchymal expression. These data show dynamic expression of Hh signaling components in the developing trachea and suggest that Hh signaling may operate in both epithelial and mesenchymal tracheal cell development.

**FIGURE 1.**
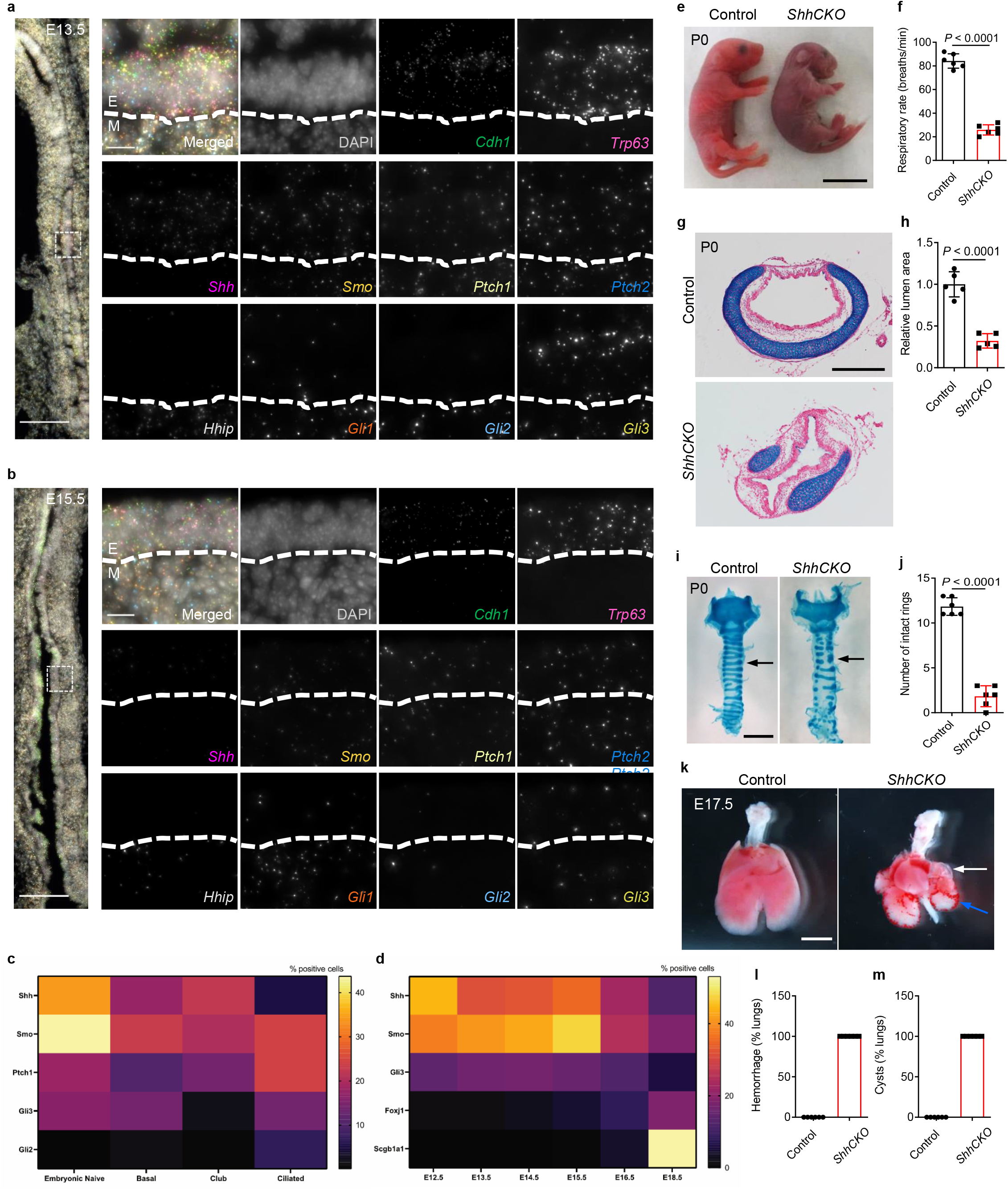
*Nkx2.1*^*Cre*^*;Shh*^*flox/flox*^ mice exhibit fracture of the tracheal cartilage, pulmonary hemorrhage and cystic lungs. (**a-b**) mRNA *in situ* detection of *Cdh1* (green), *Trp63* (wine), *Shh* (magenta), *Smo* (yellow), *Ptch1* (sand), *Ptch2* (blue), *Hhip* (pale-green), *Gli1* (orange), *Gli2* (cyan) and *Gli3* (olive) expression and DAPI staining (grey) of longitudinal sections of WT tracheae at different developmental stages. The dashed insert square in the large image indicates the area enlarged and rotated in the right side of the panels. Dashed lines in small images on the right side indicate the border between the tracheal epithelium and mesenchyme. **(c)** Quantification of the percentage of positive cells within each cell state that express Hh signaling components. **(d)** Quantification of the percentage of positive cells that express Hh signaling components. Number of cells at E12.5=239, E13.5=455, E14.5=1232, E15.5=931, E16.5=365, E18.5=286. **(e)** Representative gross morphology of P0 control (*n* = 6) and *ShhCKO* mutants (*n* = 6). (**f)** Quantification of P0 control (*n* = 6) and *ShhCKO* mutant (*n* = 6) respiratory rate. (**g**) Representative images of transverse sections of tracheae stained with alcian blue and nuclear fast red from P0 control (*n* = 5) and *ShhCKO* mutants (*n* = 5). **(h)** Quantification of P0 control (*n* = 5) and *ShhCKO* mutant (n=5) tracheal lumen area. (**i)** Representative images of ventral views of whole-mount tracheae stained with alcian blue from P0 control (*n* = 6) and *ShhCKO* mutants (*n* = 6). Arrows point to tracheal cartilage rings. (**j**) Quantification of the number of intact tracheal cartilage rings from P0 control (*n* = 6) and *ShhCKO* mutants (*n* = 6). (**k)** Representative images of ventral views of the trachea and lungs from P0 control (*n* = 6) and *ShhCKO* mutants (*n* = 6). White arrows point to pulmonary cysts. Blue arrows point to blood spots. **(l)** Percentage of lungs with hemorrhage. **(m)** Percentage of lungs with cysts. Scale bars: 2000 μm (**k**), 1000 μm (**e, i**), 400 μm (**g**), 100 μm (tracheal large images in **a, b**), 10 μm (tracheal small images in **a, b**). Unpaired Student’s *t*-test, mean ± s.d. (**f, h, j, l, m**). E, epithelium; M, Mesenchyme; Shh, Sonic Hedgehog; Smo, Smoothened; *ShhCKO*, Sonic Hedgehog conditional knockout.

To identify single embryonic cells that express Hh signaling components and relate their expression with known cell types of the trachea, we used publicly available single cell transcriptomic data, from E12.5 to E18.5 stage wild-type (WT) tracheal epithelial cells (Kiyokawa et al., 2021). We projected the expression of a panel of Hh signaling components on these data (Supplementary Fig. 3) and kept the cell type classification of the published study. We found *Shh* and *Smo* to be highly expressed in embryonic tracheal epithelial cells and the majority of the embryonic cells were positive for *Shh* or *Smo*. There were also a few embryonic cells that co-expressed both the ligand and the transducer. At later stages (E16.5 and E18.5), when known cell type markers initiate their expression, we detected the expression of *Shh* or *Smo* in a few ciliated, club and basal cells (Fig. 1c). Moreover, co-expression of *Shh* and *Smo* was only detected in rare club or basal cells but not in ciliated cells. The number of epithelial cells expressing Shh signaling components increased up to E15.5 and was reduced in later stages when cell type specific markers were expressed (Fig. 1d). Recently, *Ihh* was also reported to be expressed by alveolar epithelial cells (Nikolić et al., 2017), but at comparatively very low levels and only in a few embryonic tracheal epithelial cells (Supplementary Fig. 3d). These data suggest an autocrine function of Hh signaling on epithelial cells in early stages of trachea development (up to stage E16). We conclude that epithelial Hh signaling is activated in the trachea and hypothesize that the autocrine function of Hh signaling on the tracheal epithelium could occur in two ways, either in a cell-autonomous autocrine fashion or in a paracrine fashion between epithelial cells of different types.

*Shh* mutants exhibit defects in tracheal cartilage formation during embryonic stages (Park et al., 2010; Yin et al., 2018). We thus hypothesized that epithelial deletion of *Shh* might cause tracheomalacia with respiratory distress. To inactivate *Shh* expression specifically in the tracheal epithelium, we used a BAC transgenic mouse line where Cre recombinase is expressed under the control of *Nkx2*.*1* control elements (*Nkx2*.*1*^*Cre*^) (Xu et al., 2017). The recombination efficiency of *Nkx2*.*1*^*Cre*^ in the trachea and lungs has been previously characterized in *Nkx2*.*1*^*Cre*^*;ROSA26R-LacZ* embryos (Sala et al., 2011; Tiozzo et al., 2009). In these experiments, LacZ activity was detected in the lung epithelium as early as E10.5 and the pattern of LacZ activity was nearly homogeneous throughout the tracheal epithelium at E13.5 (Sala et al., 2011; Tiozzo et al., 2009). Thus, the *Nkx2*.*1*^*Cre*^ strain induces recombination in tracheal epithelial cells with high efficiency. We further evaluated the recombination efficiency of the *Shhflox* allele, by RT-qPCR for *Shh* mRNA expression in WT embryos and in *Nkx2*.*1*^*Cre*^*;Shh*^*flox/flox*^ (*ShhCKO*) mutants. We observed a strong reduction of *Shh* mRNA levels in both E11.5 and E18.5 stages in *ShhCKO* tracheae compared with controls (Supplementary Fig. 4). *ShhCKO* mice displayed cyanosis (Fig. 1e), neonatal respiratory distress (Fig. 1f and Supplementary Video), and died within 24 hours of birth. These mutants were born in the expected Mendelian ratio with a collapsed tracheal lumen (Fig. 1g, h), and fractured cartilage rings instead of the intact ventrolateral cartilage rings seen in control siblings (Fig. 1i, j), indicating that epithelial inactivation of *Shh* does not cause embryonic lethality. *ShhCKO* mice exhibited no trachea-esophageal fistula at E17.5. Although *Shh* mRNA levels in E11.5 *ShhCKO* lungs were already severely reduced, the lack of fistula may suggest that *Shh* inactivation in this strain occurs after the separation of the trachea from the esophagus (Supplementary Fig. 5a, b). To test for a role of *Shh* in lung development, we analyzed the embryonic lungs at E17.5. As expected, *ShhCKO* animals also exhibited pulmonary hemorrhage (Fig. 1k, l) and cysts (Fig. 1k, m), mainly in distal regions compared with controls (Miller et al.,2004). These data indicate that the mutants die from compromised respiratory function due to defective formation of the tracheal tube with high airflow resistance limiting breathing frequency and respiratory air sacs additionally impacting gas exchange.

### Epithelial cell differentiation and proliferation defects in the ShhCKO trachea

Sonic Hedgehog signaling controls epithelial differentiation in endodermal organs including the esophagus and thymus (Freestone et al., 2003; van Dop et al., 2013; Saldaña et al., 2016). To test for a possible role of *Shh* signaling in tracheal epithelial cell differentiation, we performed immunostaining for characteristic markers of major tracheal epithelial cell types. FOXJ1, a ciliated cell marker, is expressed in the trachea around E15.5 (Stauber et al., 2017). We observed several FOXJ1^+^ but no acetylated alpha-tubulin^+^ cells in the WT tracheal epithelium at E15.5 (Supplementary Fig. 6a, b). Acetylated alpha-tubulin^+^ ciliated cells were detected at E16.5 (Supplementary Fig. 6b), indicating that mature ciliated cells first appear in the trachea between E15.5 and E16.5. From E18.5, an even distribution of acetylated alpha-tubulin^+^ ciliated cells was observed in the tracheal epithelium (Supplementary Fig. 6b). Interestingly, *ShhCKO* animals at E18.5 exhibited reduced FOXJ1^+^ ciliated cells (Fig. 2a, b) with decreased *Foxj1* mRNA levels compared to controls (Fig. 2c). Consistently, the relative number of mature ciliated cells expressing acetylated alpha-tubulin (Fig. 2d, e), and the overall *Tubb4b* mRNA levels (Fig. 2c), were also reduced in *ShhCKO* animals. Next, we examined markers of club cell maturation in the mutant epithelium. SCGB1A1, a mature club cell marker, was readily detected in the WT tracheal epithelium at E18.5 (Supplementary Fig. 6c). These SCGB1A1^+^ cells were distributed evenly in the apical epithelial surface and expressed variable SCGB1A1 levels at this stage of development (Supplementary Fig. 6c). *ShhCKO* tracheal sections from E18.5 embryos exhibited fewer SCGB1A1^+^ club cells (Fig. 2f, g) and decreased *Scgb1a1* mRNA levels compared with controls (Fig. 2c). Intriguingly, the earlier secretory club cell fate markers SCGB3A2 and SCGB3A1 (Guha et al., 2012; Reynolds et al., 2002), which were detected in the WT trachea already at E17.5 (Supplementary Fig. 6d, e), exhibited a similar distribution in *ShhCKO* and WT tracheae (Supplementary Fig. 7a-d). The overall levels of *Scgb3a2* and *Scgb3a1* mRNA in the mutant trachea were also similar to controls (Supplementary Fig. 7e). These results indicate that Hh signaling is required for the progression but not the initiation of the secretory club cell differentiation program. To further examine epithelial differentiation defects in the trachea at a postnatal stage, we performed immunostaining in tissue sections at P0. *ShhCKO* tracheae exhibited persistently reduced numbers of acetylated alpha-tubulin^+^ ciliated cells (Supplementary Fig. 8a, b) and of SCGB1A1^+^ club cells (Supplementary Fig. 8a, c) compared with controls, arguing that these defects are not due to a general delay of differentiation. Since the transcription factors FOXP1 and FOXP4 cooperatively modulate club cell differentiation in the airways (Li et al., 2012), we examined for changes in *Foxp1* and *Foxp4* expression. *ShhCKO* tracheae exhibited reduced *Foxp1* expression (Fig. 2c), suggesting that *Foxp1* is a potential target of Hh signaling during club cell development.

**FIGURE 2.**
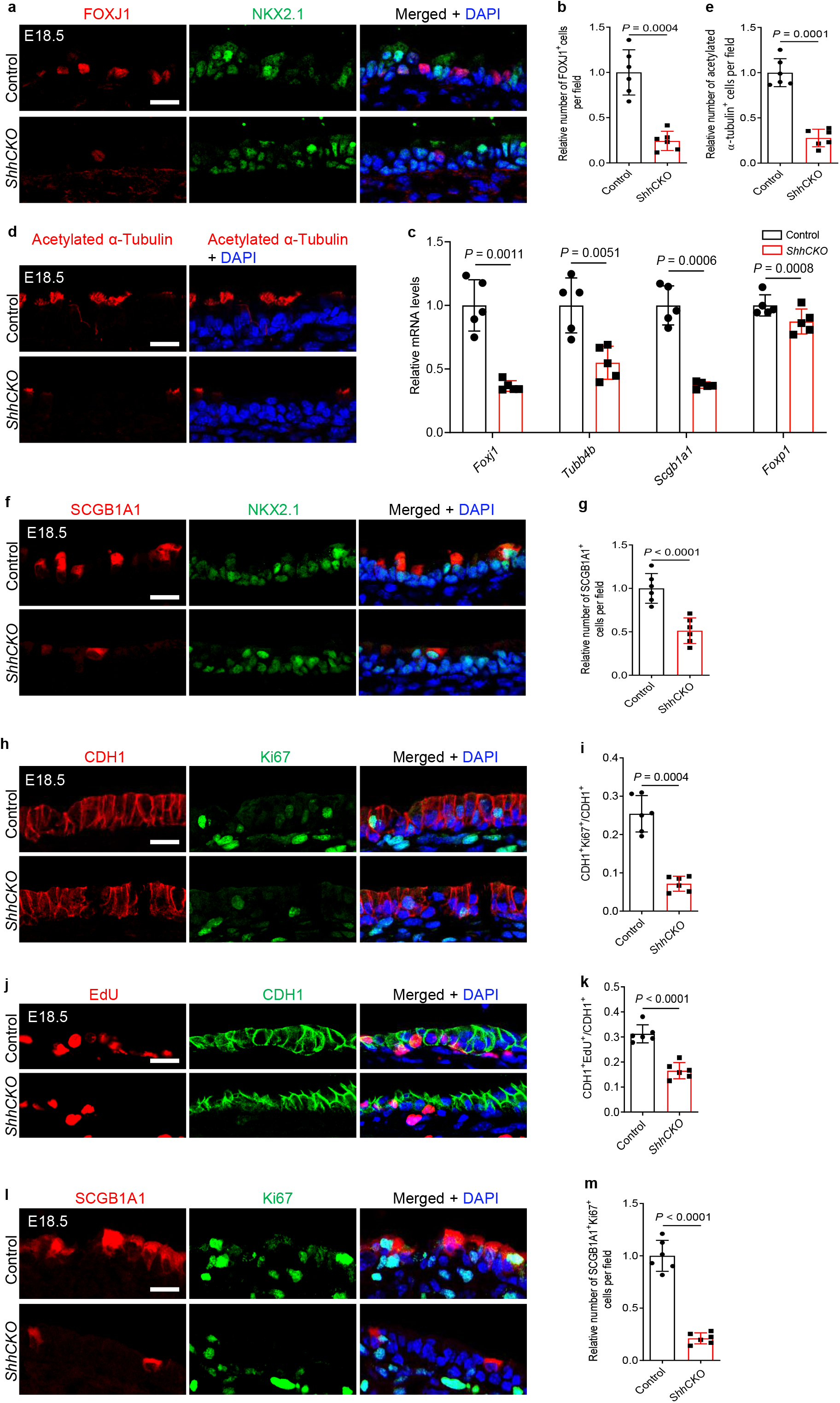
*Nkx2.1*^*Cre*^*;Shh*^*flox/flox*^ mice exhibit defects in tracheal epithelial cell proliferation and differentiation. (**a**) Immunostaining for FOXJ1 (red), NKX2.1 (green) and DAPI staining (blue) of longitudinal sections of E18.5 control (n=6) and *ShhCKO* (n=6) tracheae. (**b**) Quantification of the relative number of FOXJ1^+^ cells in the tracheal epithelium in E18.5 control (n=6) and *ShhCKO* (n=6) mice. (**c**) RT-qPCR analysis of *Foxj1, Tubb4b, Scgb1a1* and *Foxp1* mRNA levels in E18.5 control (n=5) and *ShhCKO* (*n*=5) tracheae. (**d**) Immunostaining for acetylated alpha-tubulin (red) and DAPI staining (blue) of longitudinal sections of E18.5 control (n=6) and *ShhCKO* (n=6) tracheae. (**e**) Quantification of the relative number of acetylated alpha-tubulin^+^ cells in the tracheal epithelium in E18.5 control (n=6) and *ShhCKO* (n=6) mice. (**f**) Immunostaining for SCGB1A1 (red), NKX2.1 (green) and DAPI staining (blue) of longitudinal sections of E18.5 control (n=6) and *ShhCKO* (n=6) tracheae. (**g**) Quantification of the relative number of SCGB1A1^+^ cells in the tracheal epithelium in E18.5 control (n=6) and *ShhCKO* (n=6) mice. (**h**) Immunostaining for CDH1 (red), Ki67 (green) and DAPI staining (blue) of longitudinal sections of E18.5 control (n=6) and *ShhCKO* (n=6) tracheae. (**i**) Ratio of CDH1^+^ cells that are Ki67^+^. (**j**) EdU fluorescence (red), immunostaining for CDH1 (green) and DAPI staining (blue) of longitudinal sections of E18.5 control (n=6) and *ShhCKO* (n=6) tracheae. (**k**) Ratio of CDH1^+^ cells that are EdU^+^. (**l**) Immunostaining for SCGB1A1 (red), Ki67 (green) and DAPI staining (blue) of longitudinal sections of E18.5 control (n=6) and *ShhCKO* (n=6) tracheae. (**m**) Ratio of SCGB1A1^+^ cells that are Ki67^+^. Scale bars: 20 μm. Unpaired Student’s *t*-test, mean ± s.d.

Hh signaling has been reported to modulate cell proliferation in several contexts (Berman et al., 2003; Thayer et al., 2003; van den Brink et al., 2004; Merchant et al., 2010; Plaisant et al., 2011; Gagné-Sansfaçon et al., 2014; Raleigh et al., 2018). We thus tested whether epithelial deletion of *Shh* might lead to tracheal epithelial cell proliferation defects. We examined epithelial cell proliferation first by immunostaining for Ki67, a cell cycle marker, and found that *ShhCKO* tracheae exhibited a reduced number of Ki67^+^ cells in the CDH1 marked epithelium compared with controls (Fig. 2h, i). Next, we performed an EdU incorporation assay to assess cells that had entered S-phase during the labeling period. After an *ex vivo*, 21 hours long EdU treatment, *ShhCKO* tracheae displayed fewer EdU^+^ cells in their epithelium (Fig. 2j, k), indicating that Hedgehog inactivation compromises cell cycle entry. In a more detailed analysis, we observed a reduced number of Ki67^+^SCGB1A1^+^ cells (Fig. 2l, m), but a similar number of PDPN^+^ (Supplementary Fig. 9a, b) and Ki67^+^PDPN^+^ basal cells (Supplementary Fig. 9a, c) in *ShhCKO* tracheae compared with controls. Interestingly, *ShhCKO* tracheae exhibited increased ratios of KRT5^+^ basal cells to acetylated alpha-tubulin^+^ ciliated cells and to SCGB1A1^+^ club cells compared with controls (Supplementary Fig. 10a-d). We did not find obvious alterations in Caspase3 stainings between *ShhCKO* tracheal epithelium and controls at E18.5, suggesting that apoptosis is not affected in the mutants (Supplementary Fig. 9d). To examine for changes in the surrounding mesenchyme, we analyzed SOX9^+^ chondroblasts, α-SMA^+^ smooth muscle cells and mesenchymal cell proliferation. *ShhCKO* tracheae exhibited SOX9^+^ chondroblast condensation defects (Supplementary Fig. 11a), but no obvious SOX9^+^ chondroblast differentiation defects (Supplementary Fig. 11b-d). We also detected reduced numbers of α-SMA^+^ smooth muscle cells and increased numbers of CDH1^-^Ki67^+^ mesenchymal cells compared with controls (Supplementary Fig. 11e, f, g). Since Hh signaling modulates *Fgf10* expression in the lung mesenchyme (Volckaert et al., 2013; Balasooriya et al. 2017), we examined *Fgf10* mRNA expression levels in *ShhCKO* tracheae. We detected a 1.5 fold increase in *Fgf10* mRNA in *ShhCKO* mutants (Supplementary Fig. 11h) but this was not sufficient to increase basal cell number, as might have been expected from previous studies. Altogether, these results indicate that loss of *Shh* function reduces club cell proliferation and interferes with ciliated cell differentiation and secretory cell maturation in the tracheal epithelium. In the tracheal mesenchyme, *Shh* promotes chondrocyte and smooth muscle morphogenesis.

### SMO is required in tracheal epithelial cells for cell proliferation and differentiation

To investigate whether epithelial inactivation of Hh signaling causes phenotypes similar to those observed in the *ShhCKO* tracheal epithelium, we selectively deleted the *Shh* effector *Smo* in epithelial cells by using *Nkx2*.*1*^*Cre*^*;Smo*^*flox/flox*^ (*SmoCKO*) mice. The recombination efficiency of the *Shhflox* and *Smoflox* alleles has been investigated with a *Nestin*-driven Cre transgene (*Nestin*^*Cre*^) (Machold et al. 2003). In the neural progenitor cells, the phenotypes of *Shh*^*null*^*/Shh*^*flox*^ and *Smo*^*null*^*/Smo*^*flox*^ mutant mice were indistinguishable, suggesting that the *Shhflox* and the *Smoflox* are recombined with similarly high efficiency (Machold et al. 2003). Additionally, *Act2*^*Cre*^*;Smo*^*flox*^*/Smo*^*flox*^ embryos display similar phenotypes to the *Smo*^*null*^*/Smo*^*null*^ mutants indicating that *Smo*^*flox*^ recombines with high efficiency. *ShhCKO* embryos display a strong reduction of *Shh* mRNA in their lungs, suggesting that *SmoCKO* and *ShhCKO* represent strong loss of function mutants (Supplementary Fig. 4). We detected decreased numbers of FOXJ1^+^ ciliated cells in *SmoCKO* mice (Fig. 3a, b), and an overall reduction of *Foxj1* mRNA levels (Fig. 3c) in mutant trachea compared with controls at E18.5. Similarly, we detected fewer acetylated alpha-tubulin^+^ mature ciliated cells (Fig. 3d, e) and reduced *Tubb4b* mRNA levels (Fig. 3c) in *SmoCKO* animals. *SmoCKO* tracheae also exhibited decreased numbers of SCGB1A1^+^ club cells (Fig. 3f, g) and lower *Scgb1a1* mRNA levels throughout the tissue (Fig. 3c). Interestingly, *SmoCKO* tracheae exhibited increased ratios of KRT5^+^ basal cells to acetylated alpha-tubulin^+^ ciliated cells and to SCGB1A1^+^ club cells compared with controls (Supplementary Fig. 12a-d). As in the *ShhCKO* mutants, the *SmoCKO* tracheae did not exhibit significant defects in the numbers of SCGB3A2^+^ (Supplementary Fig. 13a, b) or SCGB3A1^+^ (Supplementary Fig. 13c, d) cells, or in the overall expression levels of *Scgb3a2* or *Scgb3a1* (Supplementary Fig. 13e) compared with controls. *SmoCKO* tracheae also displayed reduced EdU^+^ incorporation in CDH1^+^ epithelial cells (Fig. 3h, i) compared with controls, suggesting that epithelial *Smo* controls cell proliferation. However, we did not detect any defects in cartilage ring formation or chondroblasts and smooth muscle cells (Supplementary Fig. 14a-e) in *SmoCKO* tracheae. Notably, the observed phenotypes in *SmoCKO* mutants are not as severe as those in *ShhCKO* animals, suggesting that the increased severity in the *ShhCKO* tracheae may be due to the reciprocal signaling from loss of *Shh* signaling in the surrounding mesenchyme. SHH activation in mesenchymal cells may also indirectly contribute to tracheal epithelial development.

**FIGURE 3.**
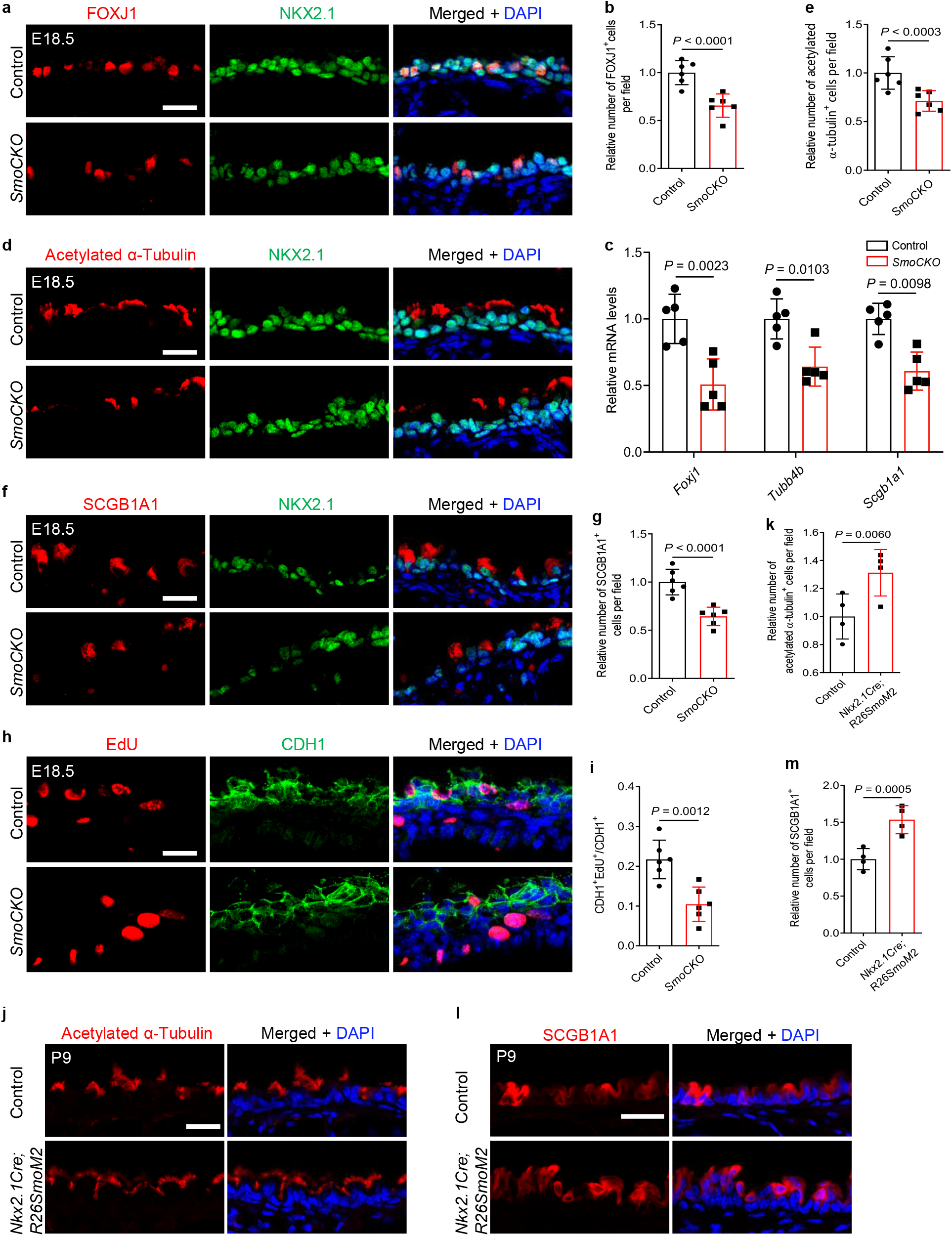
*Smo* mediates Hh-signaling in tracheal epithelial cells and controls their differentiation and proliferation. (**a**) Immunostaining for FOXJ1 (red), NKX2.1 (green) and DAPI staining (blue) of longitudinal sections of E18.5 control (n=6) and *SmoCKO* (n=6) tracheae. (**b**) Quantification of the relative number of FOXJ1^+^ cells in the tracheal epithelium in E18.5 control (n=6) and *SmoCKO* (n=6) mice. (**c**) RT-qPCR analysis of *Foxj1, Tubb4b* and *Scgb1a1* mRNA levels in E18.5 control (n=5) and *SmoCKO* (*n*=5) tracheae. (**d**) Immunostaining for acetylated alpha-tubulin (red), NKX2.1 (green) and DAPI staining (blue) of longitudinal sections of E18.5 control (n=6) and *SmoCKO* (n=6) tracheae. (**e**) Quantification of the relative number of acetylated alpha-tubulin^+^ cells in the tracheal epithelium in E18.5 control (n=6) and *SmoCKO* (n=6) mice. (**f**) Immunostaining for SCGB1A1 (red), NKX2.1 (green) and DAPI staining (blue) of longitudinal sections of E18.5 control (n=6) and *SmoCKO* (n=6) tracheae. (**g**) Quantification of the relative number of SCGB1A1^+^ cells in the tracheal epithelium in E18.5 control (n=6) and *SmoCKO* (n=6) mice. (**h**) EdU fluorescence (red), immunostaining for CDH1 (green) and DAPI staining (blue) of longitudinal sections of E18.5 control (n=6) and *SmoCKO* (n=6) tracheae after a 21 h 10 μM EdU treatment. (**i**) Ratio of CDH1^+^ cells that are EdU^+^. (**j**) Immunostaining for acetylated alpha-tubulin (red) and DAPI staining (blue) of longitudinal sections of P9 control (n=4) and *Nkx2*.*1*^*Cre*^*;R26*^*SmoM2*^(n=4) tracheae. (**k**) Quantification of the relative number of acetylated alpha-tubulin^+^ cells in the tracheal epithelium in P9 control (n=4) and *Nkx2*.*1*^*Cre*^*;R26*^*SmoM2*^(n=4) mice. (**l**) Immunostaining for SCGB1A1 (red) and DAPI staining (blue) of longitudinal sections of P9 control (n=4) and *Nkx2*.*1*^*Cre*^*;R26*^*SmoM2*^(n=4) tracheae. (**m**) Quantification of the relative number of SCGB1A1^+^ cells in the tracheal epithelium in P9 control (n=4) and *Nkx2*.*1*^*Cre*^*;R26*^*SmoM2*^(n=4) mice. Scale bars: 20 μm. Unpaired Student’s *t*-test, mean ± s.d.

To investigate whether epithelial activation of Hh signaling is sufficient to drive tracheal epithelial cell differentiation, we overexpressed a constitutively active form of *Smo, SmoM2* (Jeong et al., 2004), in epithelial cells using *Nkx2*.*1*^*Cre*^*;R26*^*SmoM2*^ mice. *Nkx2*.*1*^*Cre*^*;R26*^*SmoM2*^ tracheae displayed increased numbers of acetylated alpha-tubulin^+^ ciliated cells (Fig. 3j, k) and SCGB1A1^+^ club cells (Fig. 3l, m) compared with controls. Overactivation of *Smo* also resulted in a weak increase in the proportion of TRP63^+^KRT5^+^ basal cells in the epithelium but this difference was not statistically significant compared with controls (Supplementary Fig. 15). Collectively, these data indicate that activation of epithelial Hh signaling is required for the proliferation of a subset of tracheal epithelial cells and for the differentiation of secretory and ciliated cell types.

### Hh signaling inhibition interferes with the differentiation of human bronchial epithelial cells

The epithelial structures of the mouse trachea share similarities with human bronchioles, including the cartilaginous rings overlaid by basal progenitor cells and apical secretory and ciliated cells (Rock et al., 2010; Danopoulos et al., 2019). To examine whether Hh signaling is required for human bronchial epithelial (HBE) cell differentiation, we used primary bronchial epithelial cells cultured *in vitro* at air-liquid interface (ALI) as a model of human airway epithelial development (Schmid et al., 2017). We seeded undifferentiated HBE cells onto transwell filters and removed the medium from the upper chamber to initiate cell differentiation (defined as day 0). We then examined the expression levels of Sonic Hedgehog signaling components in HBE cells and found that *SHH, SMO, GLI1* and *GLI2* mRNA were dynamically expressed from day 0 to day 14 (Fig. 4a-d).

**FIGURE 4.**
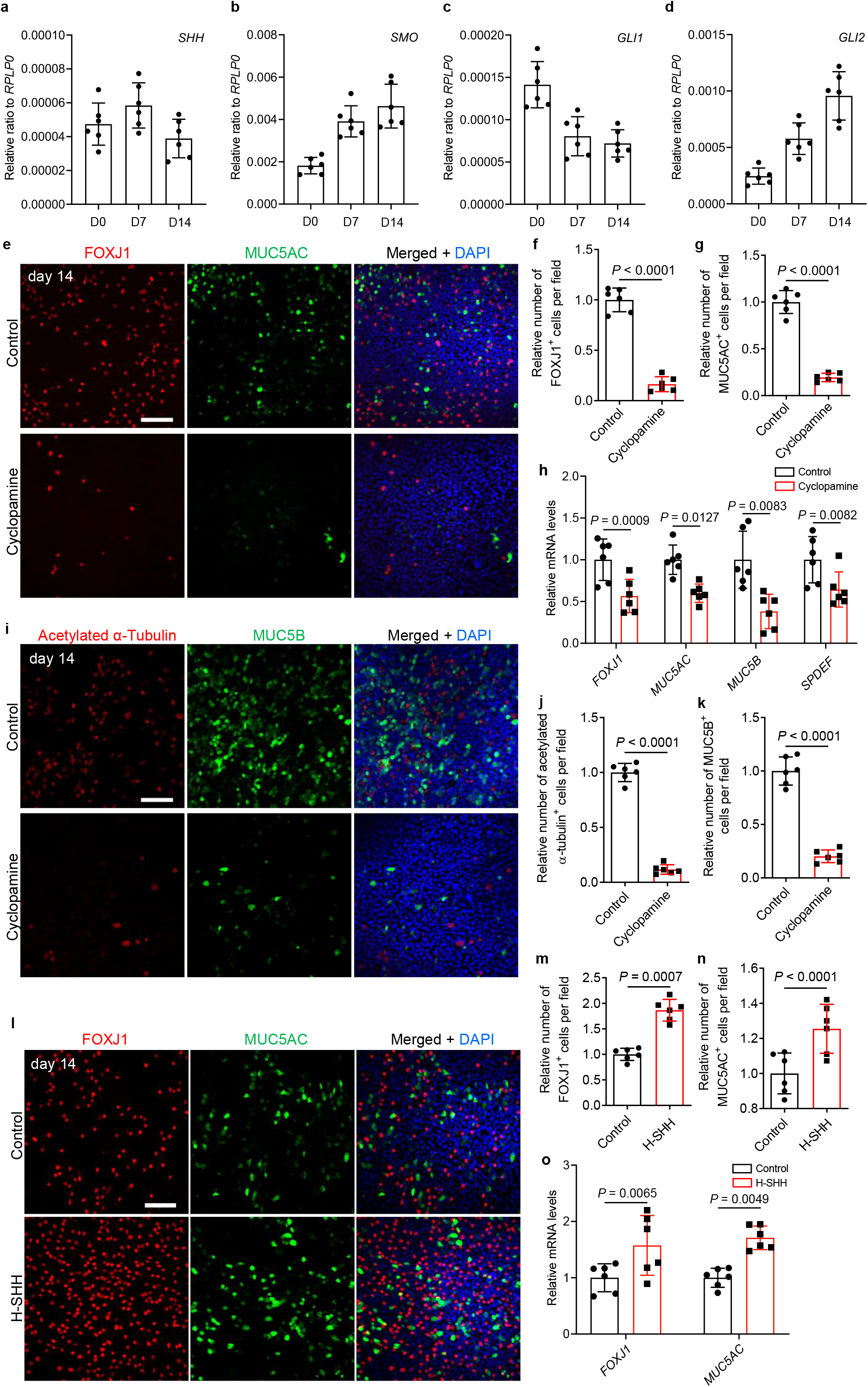
Hh signaling regulates HBE cell differentiation. **(a**-**d)** RT-qPCR analysis of mRNA levels of Hh-signaling components in ALI-cultured HBE cells (n=6 per stage). **(a)** *SHH*, **(b)** *SMO*, **(c)** *GLI1* (**d**) *GLI2*. (**e**) Immunostaining for FOXJ1 (red), MUC5AC (green) and DAPI staining (blue) in HBE cells at the ALI after 9 days of ethanol (n = 6) or 10 μM cyclopamine (n = 6) treatment. (**f**) Quantification of the relative number of FOXJ1^+^ cells in HBE cells at the ALI after 9 days of ethanol (n = 6) or 10 μM cyclopamine (n = 6) treatment. 2943 and 477 FOXJ1^+^ cells were analyzed for controls and 10 μM cyclopamine treatment respectively. (**g**) Quantification of the relative number of MUC5AC^+^ cells in HBE cells at the ALI after 9 days of ethanol (n = 6) or 10 μM cyclopamine (n = 6) treatment. 3039 and 591 MUC5AC^+^ cells were analyzed for controls and 10 μM cyclopamine treatment respectively. (**h**) RT–qPCR analysis of *FOXJ1, MUC5AC, MUC5B* and *SPDEF* mRNA levels in HBE cells at the ALI after 9 days of ethanol (n = 6) or 10 μM cyclopamine (n = 6) treatment. (**i**) Immunostaining for acetylated alpha-tubulin (red), MUC5B (green) and DAPI staining (blue) in HBE cells at the ALI after 9 days of ethanol (n = 6) or 10 μM cyclopamine (n = 6) treatment. (**j**) Quantification of the relative number of acetylated alpha-tubulin^+^ cells in HBE cells at the ALI after 9 days of ethanol (n = 6) or 10 μM cyclopamine (n = 6) treatment. 2670 and 312 acetylated alpha-tubulin^+^ cells were analyzed for controls and 10 μM cyclopamine treatment respectively. (**k**) Quantification of the relative number of MUC5B^+^ cells in HBE cells at the ALI after 9 days of ethanol (n = 6) or 10 μM cyclopamine (n = 6) treatment. 5379 and 1089 MUC5B^+^ cells were analyzed for controls and 10 μM cyclopamine treatment respectively. (**l**) Immunostaining for *FOXJ1* (red), MUC5AC (green) and DAPI staining (blue) in HBE cells at the ALI after 9 days of ddH_2_O (n = 6) or 100 ng/ml H-SHH (n = 6) treatment. (**m**) Quantification of the relative number of FOXJ1^+^ cells in HBE cells at the ALI after 9 days of ddH_2_O (n = 6) or 100 ng/ml H-SHH (n = 6) treatment. 2685 and 5007 FOXJ1^+^ cells were analyzed for controls and 100 ng/ml H-SHH treatment respectively. (**n**) Quantification of the relative number of MUC5AC^+^ cells in HBE cells at the ALI after 9 days of ddH_2_O (n = 6) or 100 ng/ml H-SHH (n = 6) treatment. 2838 and 3561 MUC5AC^+^ cells were analyzed for controls and 100 ng/ml H-SHH treatment respectively. (**o**) RT–qPCR analysis of *FOXJ1, MUC5AC* mRNA levels in HBE cells at the ALI after 9 days of ddH_2_O (n = 6) or 100 ng/ml H-SHH (n = 6) treatment. Scale bars: 100 μm. Unpaired Student’s *t*-test, mean ± s.d. HBE: human bronchial epithelial; H-SHH: recombinant human SHH.

Hh signaling inhibition by the SMO inhibitor cyclopamine (Chen et al., 2002) reduces epithelial cell differentiation in the colon (van den Brink et al., 2004), ureter (Bohnenpoll et al., 2017), and nose (Belgacemi et al., 2020). A 9-day cyclopamine treatment led to decreased differentiation of FOXJ1^+^ and RFX3^+^ ciliated cells and of MUC5AC^+^ secretory cells (Fig. 4e-g and Supplementary Fig. 16a-d), but unaltered TP63^+^ basal cell numbers (Supplementary Fig. 16a, c, e). Similarly, we detected reduced mRNA levels of *FOXJ1, TP73, MYB, MUC5AC* and *HHIP* mRNA levels but unchanged *TP63* mRNA levels compared with controls (Fig. 4h, Supplementary Fig. 17a-d). Cyclopamine treatment also caused decreased differentiation of acetylated alpha-tubulin^+^ ciliated cells and MUC5B^+^ secretory cells (Fig. 4i-k), and an overall reduction of *MUC5B* mRNA levels (Fig. 4h). Next, we examined the expression of *SPDEF*, a transcription factor gene involved in goblet cell differentiation (Park et al., 2007; Chen et al., 2009), and observed significantly reduced *SPDEF* mRNA levels after cyclopamine treatment compared with controls (Fig. 4h). To test whether the epithelial differentiation defects due to Hh signaling inhibition correlate with alterations in the Notch pathway we examined the levels of Notch signaling components. We did not detect significant changes in *JAG1, JAG2, NOTCH2, HES1* or *HEY1* mRNA levels upon cyclopamine treatment (Supplementary Fig. 17e-i). Moreover, we did not observe significant changes in the number of NGFR^+^ basal cells (Supplementary Fig. 18a, b). A 9-day treatment with 10 μM GANT 58, a GLI1 inhibitor (Lauth et al., 2007; Beauchamp et al., 2009), also decreased the relative numbers of FOXJ1^+^ ciliated cells (Supplementary Fig. 19a, b) and MUC5AC^+^ secretory cells (Supplementary Fig. 19a, c), as well as *HHIP* mRNA levels (Supplementary Fig. 19d), suggesting that Hh signaling regulates HBE cell differentiation in a GLI1-dependent manner. Next, we asked whether SHH could induce HBE cell differentiation. A 9 day-long treatment with recombinant human SHH (H-SHH) increased the relative number of FOXJ1^+^ ciliated cells (Fig. 4l, m) and MUC5AC^+^ secretory cells (Fig. 4l, n), accompanied by up-regulated *FOXJ1, MUC5AC* and *HHIP* mRNA levels (Fig. 4o and Supplementary Fig. 19e), at day 14 compared with controls. These data indicate that canonical Sonic Hedgehog signaling drives HBE cell differentiation.

### Hh signaling inhibition compromises HBE cell proliferation

To investigate Hh signaling in HBE cell proliferation, we analyzed the number of HBE cells in S-phase in ALI cultures with or without cyclopamine treatment. We conducted a dual pulse labeling using EdU and BrdU incorporation during HBE cell proliferation from day 0 to day 7 to follow proliferation at different stages of culture and differentiation (Fig. 5a). High Alexa Fluor 555-derived fluorescence intensity was observed in most HBE cells after a 12-hour EdU treatment at day 0 (Supplementary Fig. 20a, b), indicating that undifferentiated HBE cells undergo active proliferation. To examine proliferation characteristics of these EdU^+^ cells, we traced cell proliferation at later stages by adding BrdU into the culture medium from day 2.5 to day 6.5 and performing staining and analysis 12 hours after each pulse labeling (Fig. 5a). At day 3, both high and low Alexa Fluor 555-derived fluorescence intensities were observed in HBE cells (Fig. 5b), indicating different proliferative activities among HBE cells from day 0 to day 3. High Alexa Fluor 488-derived fluorescence intensity reflecting high BrdU incorporation was observed only in some of the cells with low Alexa Fluor 555-derived fluorescence intensity at day 3 (Fig. 5b), indicating further proliferation of some dividing cells from day 0. Ki67 was detected mainly in BrdU^+^ cells (Fig. 5b) and a subset of cells with low Alexa Fluor 555-derived fluorescence intensity (Fig. 5b). At day 5, cells with lower Alexa Fluor 555-derived fluorescence intensity were observed compared with day 3 (Fig. 5b, c), and BrdU^+^ or Ki67^+^ signals were detected in only a subset of these cells (Fig. 5c), indicating that few of the EdU^+^ cells divided from day 3 to day 5. At day 7, even fewer BrdU^+^ or Ki67^+^ cells exhibited low or undetectable Alexa Fluor 555-derived fluorescence intensity compared with day 5 (Fig 5d), suggesting that only a few of the EdU^+^ labelled cells at day 0 continue to divide from day 5 to day 7, or that the label became diluted after serial rounds of proliferation. Cyclopamine treatment led to a general decrease in BrdU incorporation in cells with Alexa Fluor 555-derived fluorescence intensity as well as reduced Ki67 immunostaining at day 3 (Fig. 5e-f), day 5 (Fig. 5g-h) and day 7 (Fig. 5i-j), indicating that Hh signaling is required continuously for DNA synthesis in proliferating HBE cells. To test whether Hh signaling is required for cell proliferation during differentiation of HBE cells, we treated HBE cells in ALI with 10 μM cyclopamine from day 5 to day 14, when ciliated cells and secretory cells begin to differentiate. Again, after a 9-day, cyclopamine treatment, we observed an overall decrease in proliferation of epithelial cells (Supplementary Fig. 21a, b). Altogether, these results suggest that Hh signaling is required for HBE cell proliferation at both the cell expansion and differentiation stages.

**FIGURE 5.**
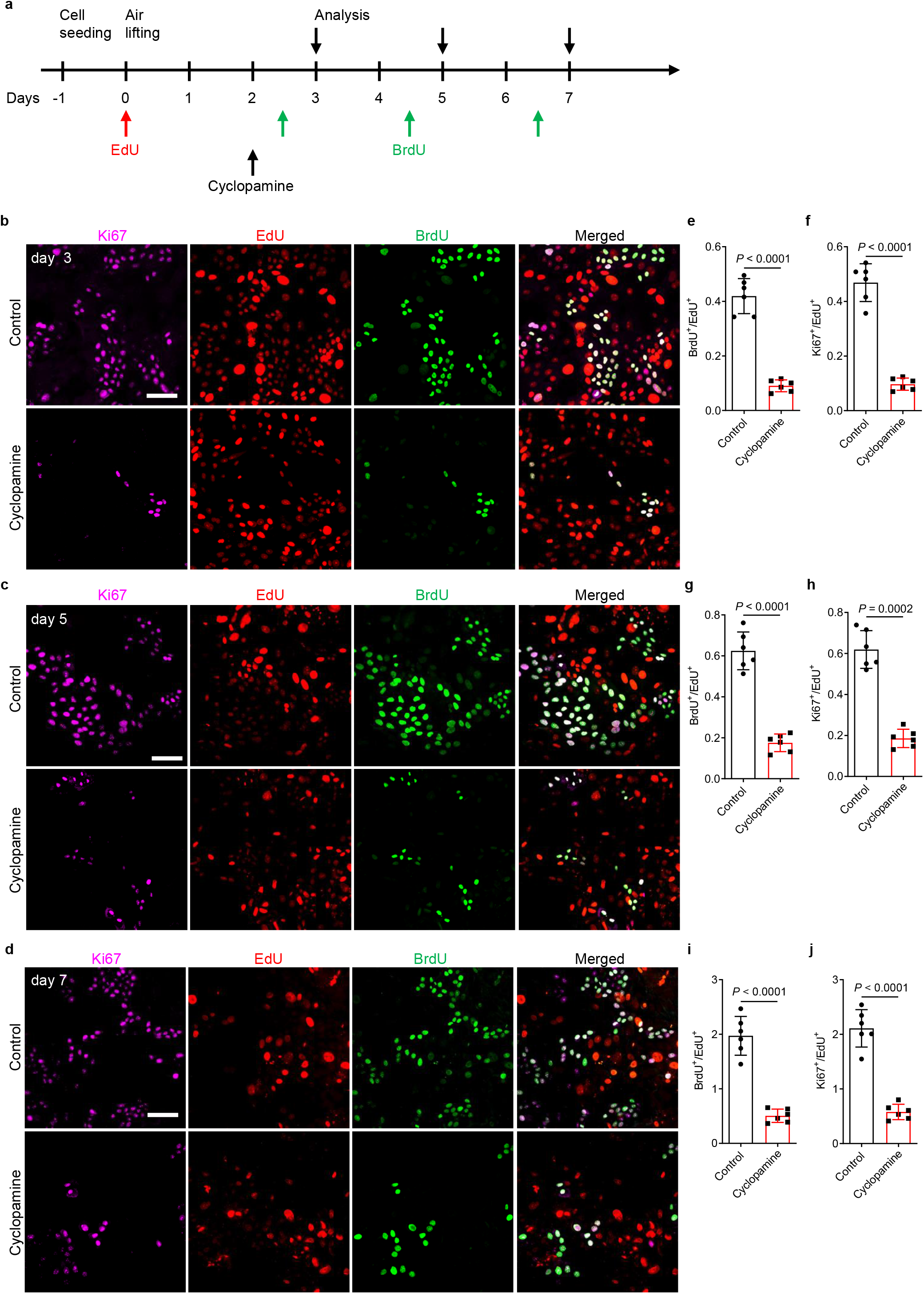
Hh signaling regulates HBE cell proliferation. (**a**) Timeline for EdU, BrdU and cyclopamine administration. (**b**) Immunostaining for Ki67 (violet), BrdU (green) and EdU fluorescence (red) in HBE cells at the ALI after ethanol (n = 6) or 10 μM cyclopamine (n = 6) treatment at day 3. (**c**) Immunostaining for Ki67 (violet), BrdU (green) and EdU fluorescence (red) in HBE cells at the ALI after ethanol (n = 6) or 10 μM cyclopamine (n = 6) treatment at day 5. (**d**) Immunostaining for Ki67 (violet), BrdU (green) and EdU fluorescence (red) in HBE cells at the ALI after ethanol (n = 6) or 10 μM cyclopamine (n = 6) treatment at day 7. (**e**) Ratio of BrdU^+^ cells to EdU^+^ cells at day 3. 1143 and 210 BrdU^+^ cells were analyzed for controls and 10 μM cyclopamine treatment respectively. (**f**) Ratio of Ki67^+^ cells to EdU^+^ cells at day 3. 1278 and 225 Ki67^+^ cells were analyzed for controls and 10 μM cyclopamine treatment respectively. (**g**) Ratio of BrdU^+^ cells to EdU^+^ cells at day 5. 1491 and 408 BrdU^+^ cells were analyzed for controls and 10 μM cyclopamine treatment respectively. (**h**) Ratio of Ki67^+^ cells to EdU^+^ cells at day 5. 1479 and 432 Ki67^+^ cells were analyzed for controls and 10 μM cyclopamine treatment respectively. (**i**) Ratio of BrdU^+^ cells to EdU^+^ cells at day 7. 1467 and 375 BrdU^+^ cells were analyzed for controls and 10 μM cyclopamine treatment. (**j**) Ratio of Ki67^+^ cells to EdU^+^ cells at day 7. 1611 and 429 Ki67^+^ cells were analyzed for controls and 10 μM cyclopamine treatment respectively. Scale bars: 100μm. Unpaired Student’s *t*-test, mean ± s.d.

## DISCUSSION

Genetic studies in mice have linked Hh signaling to several human congenital malformations of the trachea and bronchi including tracheoesophageal fistula and possible tracheomalacia (Sala et al., 2011; Miller et al., 2004; Motoyama et al., 1998; Park et al., 2009). Recently, whole-exome sequencing of pediatric patients and their parents revealed a de novo mutation in *SHH* (p.Asp279Tyr) in children with complete tracheal ring deformity (CTRD) (Sinner et al., 2019), a condition characterized by circumferentially continuous or nearly-continuous cartilaginous tracheal rings (Sahoo et al., 2009; Faust et al., 1998), variable degrees of tracheal stenosis and/or shortening, and/or pulmonary arterial sling anomaly (Berdon et al., 1984; Lee et al., 1996). Given the importance of SHH in tracheal development and congenital disorders, it is important to understand the specific contributions of paracrine and autocrine signaling activation. Our analysis of the epithelial *ShhCKO* phenotypes suggests that epithelial derived SHH mediate tracheal cartilage formation possibly by activating Hh signaling in mesenchyme. *Sox9* and *Col2a1* mRNA levels are both reduced in the tracheal mesenchyme of *Shh* null mice (Park et al., 2010), and presumably other targets involved in tracheal cartilage formation are also affected. Previous work has also shown that *Shh* and *Fgf10* exhibit genetic interaction during tracheal cartilage ring formation, and suggested that mesenchymal *Fgf10* control epithelial growth, cartilage patterning and epithelial *Shh* expression (Sala et al., 2011). Nevertheless, our epithelial cell specific deletion of *Shh* leads to more severe tracheal cartilage defects compared with the *Fgf10* or *Fgfr2b* null allele (Sala et al., 2011), suggesting that *Shh* acts via additional target (s) in the mesenchyme to regulate tracheal patterning. In bladder, epithelial cell derived SHH increases stromal expression of *Wnt2* and *Wnt4*, which in turn stimulates the proliferation of both epithelial and stromal cells on injury (Shin et al., 2011). Interestingly, both global knockout of *Wnt4* and epithelial cell specific deletion of *Wntless*, a cargo receptor facilitating Wnt ligand secretion (Bänziger et al., 2006), causes defects in tracheal cartilage ring formation (Snowball et al., 2015; Caprioli et al., 2015). Wnt signaling is also essential for human airway epithelial cell differentiation and proliferation (Schmid et al., 2017). However, it is unknown whether Wnt signaling is required for tracheal epithelial cell differentiation or proliferation *in vivo*. It would be interesting to examine epithelial cell differentiation in Wnt signaling inactivation mutants, and test for the possible crosstalk between Hh and Wnt signaling during tracheal formation.

The *SmoCKO* analysis also uncovers a new epithelial function for *Shh*, where *Smo* controls cell-autonomously the proliferation of epithelial cells and their differentiation into SCGB1A1^+^ club cells and ciliated cells. The decreased number of ciliated and SCGB1A1^+^ cells in the *SmoCKO* trachea could be due to the proliferation defect of a common progenitor or due to a continuous requirement of epithelial *Smo* for cell differentiation of the progenitors towards the ciliated and SCGB1A1^+^ lineages. This autocrine function of Hh signaling is also retained in human bronchial cells, where both SMO and GLI inhibition reduce epithelial proliferation and differentiation towards the secretory and ciliated lineages. The overexpression analysis of the activated *SmoM2* allele in the mouse and the effects of SHH addition to the human bronchial epithelial cell further argue for a direct role of Hh signaling in epithelial cell differentiation. Our work suggests a functional separation of the tracheal defects observed in mouse mutants and patients with mutations in Hh pathway genes. Cartilage abnormalities and other mesenchymal defects are due to the paracrine function of Shh, whereas tracheal lumen stenosis and respiratory dysfunctions maybe due to the combined functions of HH and its downstream effectors in both the epithelium and mesenchyme.

## MATERIAL AND METHODS

### Mice

All mouse experiments were performed under standard conditions in accordance with German federal ethical and animal welfare guidelines (Ethical Permit number GI 20/10, Nr. G 29/2019) and Swedish ethical regulations approved by the Northern Stockholm Animal Ethics Committee (Ethical Permit numbers N254/2014 and 15196/2018). All breeding colonies were maintained under cycles of 12-hour light and 12-hour dark. *Nkx2*.*1*^*Cre*^ (2), *Shh*^*flox*^ (2), *Smo*^*flox*^ (60), *R26Smo*^*M2*^ (41) alleles have been previously described.

### SCRINSHOT detection of mRNA

E13.5 and E15.5 WT tracheae were fixed in 4% PFA for 30-45 min at 4 °C. Longitudinal cryosections of the trachea (10 μm) were prepared and the SCRINSHOT protocol (25) was used for probe design and *in situ* detection of mRNA transcripts. Sequences of probes are listed in the supplementary table.

### Alcian blue staining of cartilage

Tracheal cryosections (10 µm) were fixed in 4% paraformaldehyde for 20 min, treated with 3% acetic acid solution for 3 min, stained in 0.05% alcian blue for 10 min and counterstained with 0.1% nuclear fast red solution for 5 min. For whole-mount staining of tracheal cartilage, dissected tracheae were fixed in 95% ethanol for 12 h followed by overnight staining with 0.03% alcian blue dissolved in 80% ethanol and 20% acetic acid. Samples were cleared in 2% KOH.

### Immunostaining of cryosections

Tracheae were dissected in PBS, fixed in 4% paraformaldehyde overnight at 4 °C, incubated in 10% sucrose and 30% sucrose for 24 h each at 4 °C, mounted in OCT embedding compound, and sectioned at 10 µm. To perform immunostaining, sections were fixed in 4% paraformaldehyde 10 min at 4 °C, followed by incubation in permeabilization solution (0.3% Triton X-100/PBS) for 15 min at RT, incubated in blocking solution (5% FBS/PBS/3% BSA) for 1 h at RT, incubated in primary antibodies overnight at 4 °C, washed, incubated in secondary antibodies for 2 h at RT, washed, and then mounted for imaging.

### Explant culture of mouse embryonic tracheae and lungs, and EdU incorporation and detection

Tracheae and lungs were isolated from E18.5 embryos and cultured using an established protocol (61). For EdU incorporation, isolated tracheae and lungs were cultured in DMEM/F-12 medium containing 10 µM EdU at 37 °C in a 5% CO_2_ incubator for 20 h. 0.1% DMSO was used as a control. Tracheae were fixed in 4% paraformaldehyde overnight at 4 °C, incubated in 10% sucrose and 30% sucrose for 24 h each at 4 °C, mounted in OCT embedding compound, and sectioned at 10 µm. To perform EdU detection and CDH1 immunostaining, sections were fixed in 4% paraformaldehyde 10 min at 4 °C followed by incubation in permeabilization solution (0.3% Triton X-100/PBS) for 15 min and in blocking solution (5% FBS/PBS/3% BSA) for 1 h at RT. Samples were incubated in EdU reaction cocktail prepared according to the manufacturer’s instruction (Click-iT® EdU Imaging Kit, Thermo Fisher Scientific, C10338) for 30 min, washed, incubated in the CDH1 primary antibody for 4 h at RT, washed, incubated in the secondary antibody for 2 h at RT, washed, and then mounted for imaging.

### HBE cell ALI culture, and chemical treatment

Primary human bronchial epithelial (HBE) cells (material obtained from the UGMLC/DZL-Biobank, Dr. C. Ruppert/ Prof. A. Günther and approved by the local ethic committee, AZ 85/15) were isolated and cultivated under air-liquid interface conditions to form well-differentiated, pseudostratified cultures as described previously (62). Isolated HBE cells were maintained and expanded (one passage) in T75 flasks in hormone- and growth factor-supplemented airway epithelial cell growth medium (AEGM, ready-to-use; PromoCell) at 37 °C in a 5% CO_2_ incubator. At 80% confluence, cells were detached with 0.05% trypsin-EDTA (Gibco) and seeded on membrane supports (12 mm Transwell culture inserts, 0.4 µm pore size, Costar) coated with 0.05 mg collagen from calf skin (Sigma-Aldrich) in ready-to-use AEGM supplemented with 1% penicillin/streptomycin. HBE cells were cultured for two days until they reached complete confluence. The apical medium was then removed and the basal medium was replaced by a 1:1 mixture of DMEM (Sigma) and ready-to-use AEGM supplemented with 60 ng/ml retinoic acid (Sigma). Cultures were maintained under air-liquid interface conditions by changing the medium in the basal filter chamber three times a week. For cyclopamine (1623, TOCRIS) treatment, a 5 mM stock solution was diluted to 10 µM. For GANT 58 (3889, TOCRIS) treatment, a 10 mM stock solution was diluted to 10 µM. For recombinant human SHH (H-SHH) (R&D Systems, 1845-SH-025) treatment, a 200 µg/µl stock solution was diluted to 100 ng/ml. Epithelial cells were cultured in differentiation medium containing the above chemicals or H-SHH at 37 °C in a 5% CO_2_ incubator from day 5 to day 14. The medium was replaced every 24 h before collection for analysis.

### EdU and BrdU incorporation and detection in HBE cells

For EdU incorporation in HBE cells, a 10 mM EdU stock solution was diluted to 10 µM. For BrdU incorporation, a 10 mM BrdU stock solution was diluted to 10 µM. HBE cells were cultured in differentiation medium with EdU or BrdU at 37 °C in a 5% CO_2_ incubator for 12 h at indicated time point. For EdU and BrdU detection, ALI cultures were fixed in 4% paraformaldehyde 20 min at RT, followed by incubation in permeabilization solution (0.3% Triton X-100/PBS) for 15 min and in blocking solution (5% FBS/PBS/3% BSA) for 1 h at RT. Samples were incubated in EdU reaction cocktail prepared accouding to the manufacturer’s instruction (Click-iT® EdU Imaging Kit, Thermo Fisher Scientific, C10338) for 30 min, washed, incubated in the BrdU antibody for 4 h at RT, washed, incubated in the secondary antibody for 2 h at RT, washed, and then mounted for imaging.

### Reverse transcription quantitative PCR (RT-qPCR)

Total RNA extraction was conducted using a miRNeasy Mini Kit (Qiagen, 217004). cDNA was synthesized using the Maxima First Strand cDNA Synthesis Kit (Thermo Fisher Scientific, K1641), according to manufacturer’s instructions. Quantitative real-time PCR was performed using LightCycler^®^ 480 II (Roche) and iTaq™ Universal SYBR^®^ Green Supermix (Bio-Rad, 1725122). The following primers were used: *mActb* forward 5′-CGGCCAGGTCATCACTATTGGCAAC-3′ and *mActb* reverse 5′-GCCACAGGATTCCATACCCAAGAAG-3′; *mShh* forward 5′-GGCTGATGACTCAGAGGTGCAAAG-3′ and *mShh* reverse 5′-GCTCGACCCTCATAGTGTAGAGAC-3′; *mGli1* forward 5′-GCCTGGAGAACCTTAGGCTGGA-3′ and *mGli1* reverse 5′-ACAGGTGCGCCAGCGTG-3′; *mGli2* forward 5′-GCCCTGGAGAGTCACCCTT-3′ and *mGli2* reverse 5′-TGCACAGACCGGAGGTAGT-3′; *mSmo* forward 5′-GAGCGTAGCTTCCGGGACTA-3′ and *mSmo* reverse 5′-CTGGGCCGATTCTTGATCTCA-3′; *mFoxj1* forward 5′-GAGTGAGGGCAAGAGACTGG-3′ and *mFoxj1* reverse 5′-TCAAGTCAGGCTGGAAGGTT-3′; *mTubb4b* forward 5′-AACCCGGCACCATGGACTCTGT-3′ and *mTubb4b* reverse 5′-TGCCTGCTCCGGATTGACCAAATA-3′; *mScgb1a1*forward 5′-ATGAAGATCGCCATCACAATCAC-3′ and *mScgb1a1* reverse 5′-GGATGCCACATAACCAGACTCT-3′; *mScgb3a1* forward 5′-ACCACCACCTTTCTAGTGCTC-3′ and *mScgb3a1* reverse 5′-GGCTTAATGGTAGGCTAGGCA-3′; *mScgb3a2* forward 5′-GCTGGTATCTATCTTTCTGCTGGTG-3′ and *mScgb3a2* reverse 5′-ACAACAGGGAGACGGTTGATGAGA-3′; *mFoxp1* forward 5′-CGAATGTTTGCTTACTTCCGACGC-3′ and *mFoxp1* reverse 5′-ACTTCATCCACTGTCCATACTGCC-3′; *hRPLRO* forward 5′-CCAGCAGGTGTTCGACAAT-3′ and *hRPLRO* reverse 5′-CAGGAAGCGAGAATGCAGA-3′; *hSHH* forward 5′-AAGGACAAGTTGAACGCTTTGG-3′ and *hSHH* reverse 5′-TCGGTCACCCGCAGTTTC-3′; *hGLI1* forward 5′-TCTCAAAGTGGGAGGCACAA-3′ and *hGLI1* reverse 5′-CCCTTAGGAAATGCGATCTG-3′; *hGLI2* forward 5′-TTATGGGCATCCTCTCTGGT-3′ and *hGLI2* reverse 5′-CGGAGCAGAGTATCCAGTAT-3′; *hSMO* forward 5′-GAAGTGCCCTTGGTTCGGA-3′ and *hSMO* reverse 5′-GCAGGGTAGCGATTCGAGTT-3′; *hFOXJ1* forward 5′-GCCTCCCTACTCGTATGCCA-3′ and *hFOXJ1* reverse 5′-GCCGACAGGGTGATCTTGG-3′; *hMUC5AC* forward 5′-GCTCAGCTGTTCTCTGGATGAG-3′ and *hMUC5AC* reverse 5′-TTACTGGAAAGGCCCAAGCA-3′; *hMUC5B* forward 5′-ACATGTGTACCTGCCTCTCTGG-3′ and *hMUC5B* reverse 5′-TCTGCTGAGTACTTGGACGCTC-3′; *hNOTCH2* forward 5′-TGGTGGCAGAACTGATCAAC-3′ and *hNOTCH2* reverse 5′-CTGCCCAGTGAAGAGCAGAT-3′; *hJAG1* forward 5′-GAATGGCAACAAAACTTGCAT-3′ and *hJAG1* reverse 5′-AGCCTTGTCGGCAAATAGC -3′; *hJAG2* forward 5′-GAGCTCTGCGACACCAATC-3′ and *hJAG2* reverse 5′-TCATTGACCAGGTCGTAGCA -3′; *hHES1* forward 5′-TTACGGCGGACTCCATGT-3′ and *hHES1* reverse 5′-AGAGGTGGGTTGGGGAGT-3′; *hHEY1* forward 5′-GATGATCAGCTTTATCCAAGAAAGA-3′ and *hHEY1* reverse 5′-CAGTTTGTACATTCACCTTTCTGC-3′; *hMYB* forward 5′-CCGGGAAGAGGATGAAAAAC -3′ and *hMYB* reverse 5′-TTTCCAGTCATCTGTTCCATTC-3′; *hTP73* forward 5′-CCACTGGTGGACTCCTATCG-3′ and *hTP73* reverse 5′-CTGTAGGTGACTCGGCCTCT-3′; *hTP63* forward 5′-GAAGATCAAAGAGTCCCTGGAA-3′ and *hTP63* reverse 5′-GCTGTTGCCTGTACGTTTCA-3′

### Antibodies

The following antibodies were used: Rat anti-CDH1 (1:200, Santa Cruz, sc-59778); Mouse anti-CDH1 (1:100, BD Biosciences, 560062); Rabbit anti Cleaved Caspase-3 (1:600, Cell Signaling Technologies, #9661); Goat anti-SCGB1A1 (1:200, Santa Cruz, T-18); Rabbit anti-NKX2.1 (1:400, Santa Cruz, H-190); Mouse anti-acetyl-alpha tubulin (1:2000, Sigma, MABT868); Mouse anti-FOXJ1 (1:400, Thermo Fisher Scientific, 14-9965-82); Rabbit anti-MUC5AC (1:400, Santa Cruz, H-160); Rabbit anti-MUC5B (1:400, Novus Biologicals, NBP1-92151); Mouse anti-BrdU (1:400, Thermo Fisher Scientific, B35141); Hamster anti-PDPN (1:20, DSHB, 8.1.1); Rat anti-SCGB3A1 (1:200, R&D Systems, MAB2954); Rat anti-SCGB3A2 (1:200, R&D Systems, MAB3465); Rabbit anti-Ki67 (1:400, Thermo Fisher Scientific, PA5-19462); Goat anti-NGFR (1:200, Santa Cruz, C-20); Mouse anti-α-SMA-Cy3 (1:1000, Sigma-Aldrich, C6198); Rabbit anti-SOX9 (1:400, Millipore, AB5535MA); Goat anti-SOX9 (1:500, R&D Systems, AF3075); Rat anti-Cd140a (1:100, Biolegend, 135901); Rabbit anti-RFX3 (1:500, Sigma, HPA035689); Mouse anti-TRP63 (1:500, Abcam, ab735); Rabbit anti-TRP63 (1:100, Cell Signaling Technology, #13109S). Chicken anti-KRT5 (1:400, Biosite, 905901).

### Imaging

Imaging of wholemount tracheae, trachea sections and ALI cultures was performed using a Nikon SMZ25, Zeiss 880 upright laser scanning confocal microscope, Zeiss 780 laser scanning confocal microscope or Zeiss Axio Observer Z.2 fluorescent microscope (Carl Zeiss Microscopy GmbH). SCRINSHOT imaging was performed using a Zeiss Axio Observer Z.2 fluorescent microscope (Carl Zeiss Microscopy GmbH) with a Colibri led light source, equipped with a Zeiss AxioCam 506 Mono digital camera and an automated stage. Quantification of cell number was performed using ImageJ (http://rsbweb.nih.gov/ij/).

### Statistical analysis

Statistical analyses were performed using GraphPad software. P values were calculated by Student’s t-test P values were calculated by Student’s t-test. P<0.05 was considered significant.

## ACKNOWLEDGEMENTS

We thank, Radhan Ramadass, Anoop Cherian and Yu Hsuan Carol Yang for imaging assistance, Ramesh-Kumar Krishnan, Thomas Sontag, Sigrid Einemann for discussions, and/or assistance.

## AUTHOR CONTRIBUTIONS

W.Y. and C.S. conceived the project, designed experiments and analyzed data; W.Y., A.L., L.M., J.K., M.G., X.L., C.L., H.W. and A.F. contributed to experiments and data analysis; J.K., C.R., A.G. prepared primary human bronchial epithelial (HBE) cells; A.S. contributed to data analysis. C.S., W.S. and D.Y.R.S. provided infrastructure and contributed to data analysis; W.Y., A.L. and C.S. wrote the manuscript. All authors commented on the manuscript. Funding for this study was provided by the National Natural Science Foundation of China (81970019), Open Project of State Key Laboratory of Respiratory Disease (SKLRD-OP-202110), Guangdong Key Research and Development Project (2020B1111330001), and ZHONGNANSHAN MEDICAL FOUNDATION OF GUANGDONG PROVINCE (ZNSA-2020001) to W.Y., the German Research Foundation (DFG), grant KFO309 (project number 284237345) and the Swedish Cancer Society to C.S. and the Max Planck Society to D.Y.R.S and C.S.

## COMPETING INTERESTS

The authors declare no competing interests.

## Notes

### Competing Interest Statement

The authors have declared no competing interest.

